# Minimal system for assembly of SARS-CoV-2 virus like particles

**DOI:** 10.1101/2020.06.01.128058

**Authors:** Heather Swann, Abhimanyu Sharma, Benjamin Preece, Abby Peterson, Crystal Eldridge, David M. Belnap, Michael Vershinin, Saveez Saffarian

## Abstract

SARS-CoV-2 virus is the causative agent of COVID-19. Here we demonstrate that non-infectious SARS-CoV-2 virus like particles (VLPs) can be assembled by co-expressing the viral proteins S, M and E in mammalian cells. The assembled SARS-CoV-2 VLPs possess S protein spikes on particle exterior, making them ideal for vaccine development. The particles range in shape from spherical to elongated with a characteristic size of 129 ± 32 nm. We further show that SARS-CoV-2 VLPs dried in ambient conditions can retain their structural integrity upon repeated scans with Atomic Force Microscopy up to a peak force of 1 nN.

## Main

COVID-19 is a pandemic disease caused by infection of SARS-CoV-2 virus^1^. With more than 5 million cases confirmed and a death toll exceeding several hundred thousand individuals, a search for antiviral therapies as well as vaccine candidates is of utmost urgency. Non-infectious virus like particles (VLPs) displaying essential viral proteins can be used to study the structural properties of the SARS-CoV-2 virions and due to their maximum immunogenicity are also vaccine candidates^2, 3^. VLPs are released from cells with similar mechanisms as fully infectious virions and resemble the shape and composition of fully infectious virions^4^. Most Coronaviruses are pathogens of zoonotic nature with different viruses infecting avian (IBV), bovine (BCoV), porcine (TGEV), feline (FIPV) and murine(MHV)^4^ species. There is evidence that Bat-SARS-CoV is the origin of the SARS-CoV virus which first appeared in human hosts in 2003^5^. The genome of SARS-CoV-2 has ~90% similarity to genome of coronaviruses previously identified in bat populations in China^1, 6^.

Expression of E and M proteins of TGEV result in release of VLPs with similar sizes to wild type TGEV^7^. Similar results are reported for IBV^8^. Similarly, expression of M, E and S proteins are shown to result in release of morphologically identical particles to wild type SARS-CoV virus^9, 10^. It is known that TGEV VLPs can induce an IFNα response in their hosts^7^. More recently human MERS-CoV VLPs were also shown to elicit an immune response and serve as vaccine candidates^11^.

Given the similarities between SARS-CoV-2 and SARS-CoV viruses^1, 12^, we set out to create SARS-CoV-2 VLPs by expressing S, M and E proteins of SARS-CoV-2 in mammalian cells. We then tested the structural integrity of the SARS-CoV-2 VLPs attached to dry glass using Atomic Force Microscopy (AFM), since SARS-CoV-2 virions have been reported to survive on solid surfaces in dry conditions for many hours^13^.

## Methods

Plasmid preparation and VLP harvest. SARS-CoV-2 M, S and E protein genes were identified from the full genome sequence of the virus^1^, these genes were then humanized and inserted in CMV driven mammalian expression vectors (see supplement for complete plasmid sequences). 24 hours after transfecting a monolayer of 293T cells with a cocktail of S, M and E plasmids, VLPs were harvested from the supernatant (see supplement for further details). The supernatant was filtered using 0.2um filter and VLPs were captured in a 40-10% sucrose step gradient and concentrated using AMICON spin filters (UFC901008, EMD Millipore, Burlington, MA). Total protein yield for purified VLP stock varied somewhat but was typically above 5 mg/mL. The purified VLPs were stable for at least a week when stored at ~ 4 °C.

Immunogold electron microscopy. VLPs were incubated with 1:1000 dilution of anti-S antibody (clone 1A9, GTX632604, GeneTex, Irvine, CA) and 1:5 dilution of goat anti-mouse IgG nanogold conjugates (BBI solutions, Crumlin, UK, distributed by TedPella, Redding, CA, cat # 15753). Negative stain electron microscopy was performed on VLPs by applying 3.5 uL of VLPs to a glow-discharged formvar-carbon coated EM grid (Ted Pella, Redding, CA) followed by two de-ionized water washes and staining with 1% uranyl acetate. Imaging was performed in a JOEL JEM1400-Plus microscope with an accelerating voltage of 120 keV.

Western Blot analysis. After triple washing the cells with PBS, 293T cells were suspended in 100ul of RIPA buffer (sc-24948, Santa Cruz, Dallas, Texas). 10μl of VLPs or cell extracts were then denatured by Laemmli sample buffer (BioRad, Hercules, CA) with 5% BME and boiled at 95C for 10min. The proteins were separated by SDS-PAGE and then transferred to a PVDF membrane (Millipore, Burlington, Massachusetts). Membranes were stained with 1:1000 dilution of Anti-SARS-CoV SΔ10 antibody ([1A9], GenTex, Irvine, CA) along with 1:1000 dilution of Anti-Membrane Protein (2019-nCoV) Polyclonal Antibody (NCV-M-005, eEnzyme, Gaithersburg, MD) and then immunoprobed with appropriate infrared secondary antibody. Both antibodies were confirmed to be specific to intended antigens (Fig. S3) Membrane was scanned with the Odyssey infrared imaging system (LI-COR, Lincoln, NE) according to the manufacturer’s manual instruction at 700nm and 800nm.

AFM sample prep and experiments. AFM experiments were performed using Dimension Icon AFM with an MLCT-BIO-DC probe (Bruker, Santa Barbara, CA, USA) in PeakForce QNM in air mode at room temperature. Glass coverslips were functionalized with anti-S antibody as follows. Glass was first cleaned and functionalized with biotin-PEG-silane as previously described^15^. The surfaces were then incubated with neutravidin (Thermo Scientific Pierce Protein Biology, Waltham, MA, USA) followed by a brief dd-H20 wash to get rid of excess neutravidin and incubation with biotinylated anti-S antibody (11-2001-B, Abeomics, San Diego, CA, USA). Surfaces thus prepared were generally devoid of debris but sometimes had step-edge character suggesting that antibody coating was less than a full monolayer (Figure S4). The VLPs suspended in assay buffer were then incubated with the functionalized surface and finally washed away via brief buffer exchange with dd-H20 followed by drying under Nitrogen gas flow. All incubations lasted 30 minutes at room temperature. Assay buffer: PBS, pH 7.

## Results

VLP purifications were first assessed biochemically. Figure 1 (see also Figure S1) shows western blot analysis of cell extracts as well as of purified VLPs. The isolated VLPs have both M and S proteins as identified in western blots. S protein appears more highly enriched in cells vs VLPs suggesting that not all expressed S protein is released on VLPs. The isolated VLPs were further analyzed by running a 10-40% sucrose gradient and sequential fractionation. VLPs were found to be within the density of 20-25% sucrose (Figure S2). S protein bands show a cleavage product which runs at a lower molecular weight from full S protein as shown in Figure 1 and more clearly visible in Figure S2. In addition both S protein as well as M protein form complexes which survive during western blot analysis and run at considerably higher molecular weight; successive dilutions results in breakup of these higher molecular weight fractions to respective monomeric bands for each protein (Figure S3).

**Figure 1.**
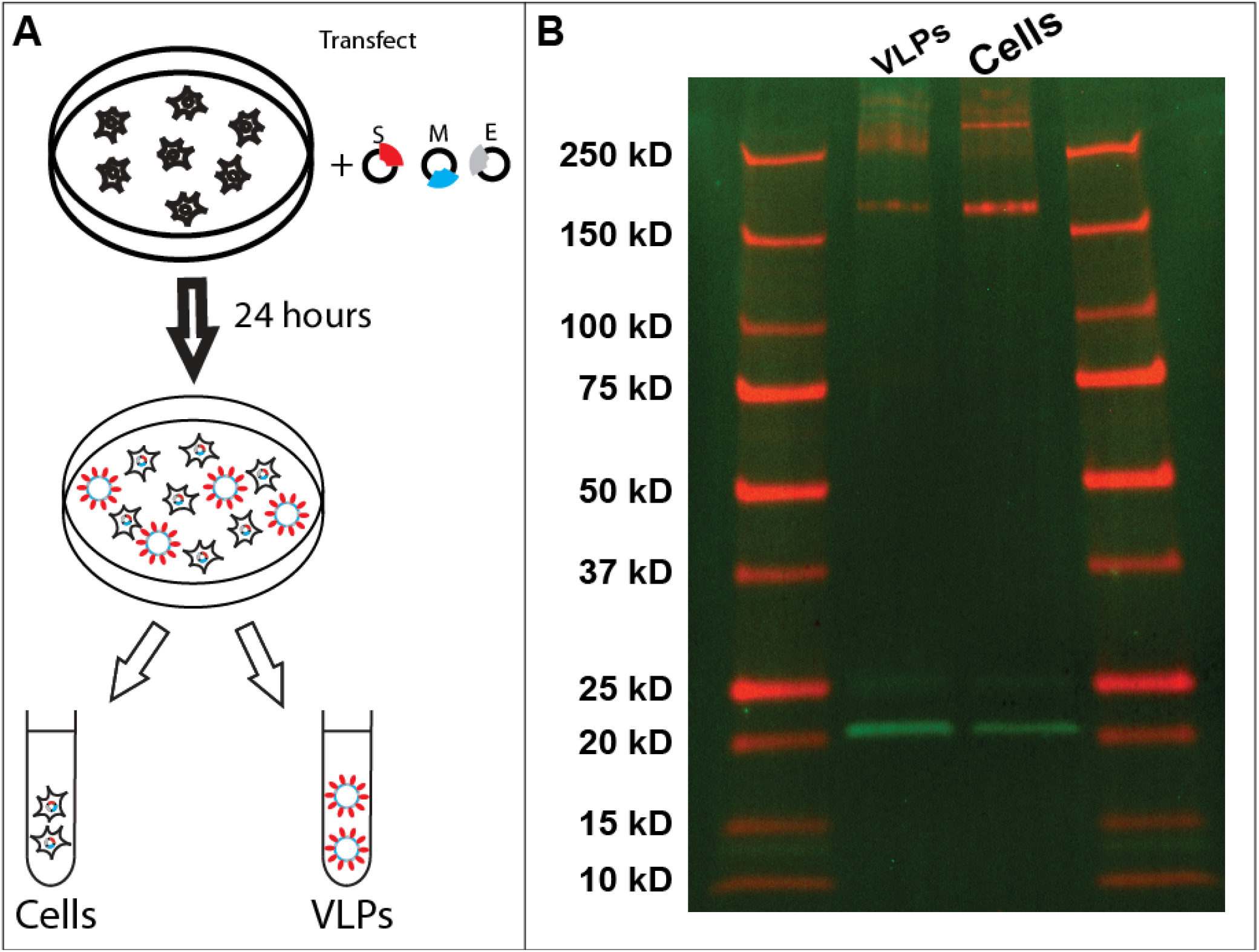
VLP expression and characterization using western blots. (a) Schematic representation of transfection and VLP harvest. (b) Western blot analysis of cells as well as VLPs separated as explained in methods. The gel was performed simultaneously, scanned at 700 nm and 800 nm, then superimposed using ImageJ (NIH) color merge for greater clarity, followed by cropping and minimal contrast enhancement. S protein (red) and M protein (green) are seen in two middle lanes. S protein tends to aggregate at very high concentrations (Fig. S2) resulting in smeared signal above 250 kD. S protein cleavage product is seen at 75 kD however the band is better visible when imaging channels are shown separately (Fig. S6). Protein size standards are described in Methods and are seen in leftmost and rightmost lanes. All size standard bands are also annotated on the left.

We further characterized the purified VLPs via electron microscopy. Figure 2 shows a representative image of the electron micrographs with the immobilized VLPs on the EM grids. VLPs can be identified via specific nanogold immunolabeling. Notably, nanogold labels not only decorate the VLPs but also proximal areas, suggestive of S protein dissociating from VLPs during surface deposition. The SARS-CoV-2 VLPs we have characterized have a size distribution of 129 ± 32 nm, consistent with the general characterization of prior coronaviruses with size distributions of 100 - 200 nanometers^4^. This is also consistent with prior reports for SARS-CoV VLPs, which were characterized with cryotomography and have size distribution of 79 – 224 nm (shape reportedly varied from nearly spherical to ellipsoidal with a 2 to 1 aspect ratio^9^). Our measured VLPs are also consistent with limited electron microscopy data available for the fully infectious SARS-CoV-2 virions observed in patient samples^14^.

**Figure 2.**
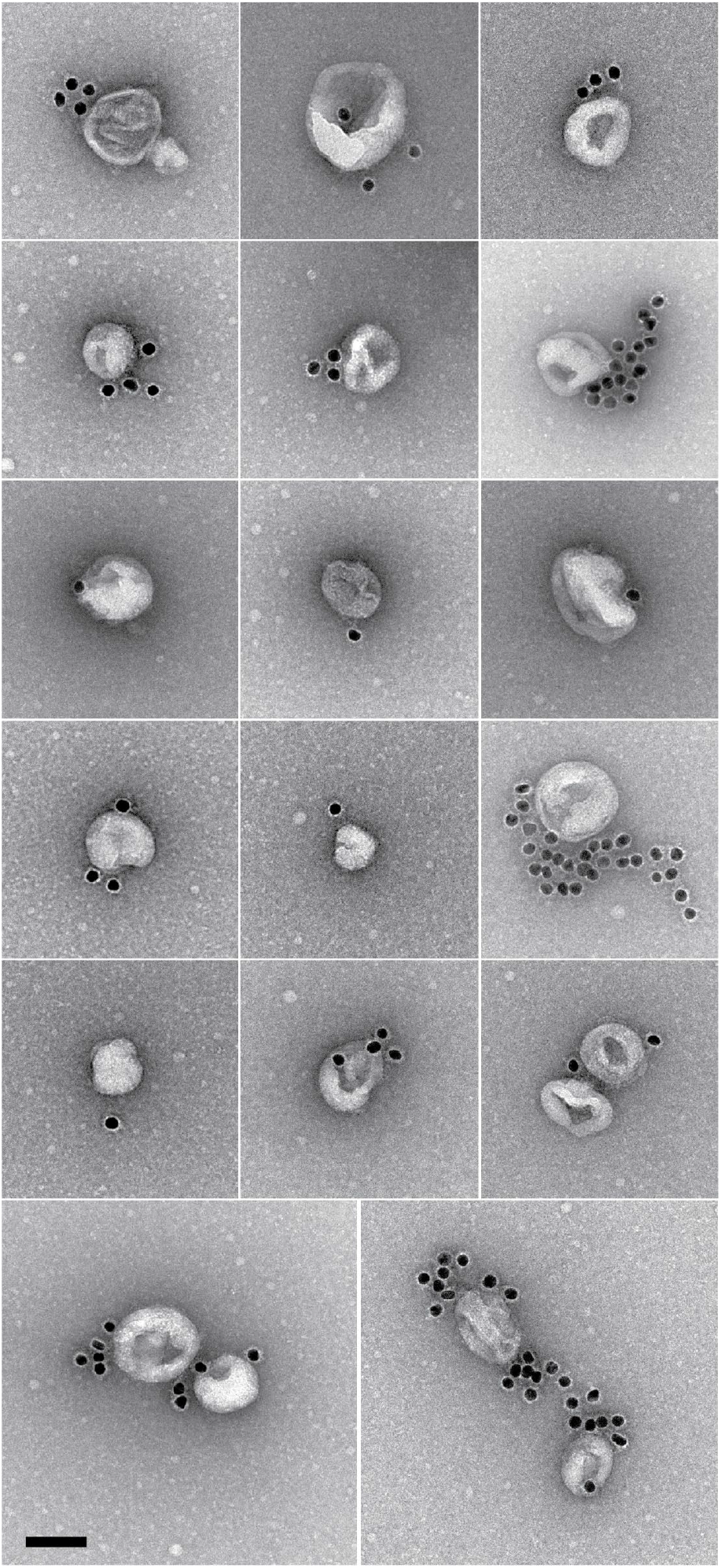
Negative stain electron microscopy reveals SARS-CoV-2 VLPs specifically labeled with immunogold. Scale bar: 100 nm.

The structural integrity of VLPs can inform their practical applications as well as serve as an estimate of the stability of fully infectious SARS-CoV-2 virions. We therefore further characterized the VLPs on functionalized surfaces with atomic force microscopy (AFM) (Fig. 3 and Figure S5 & S6). VLPs were independently imaged on multiple occasions, originating from multiple purification batches. The observed surface density was as high as ~100 per 10 μm square field of view for high-yield VLP batches. VLPs appeared as approximately spherically symmetric particles whose lateral diameter was in excess of 200 nm while heights of the particles varied between 50 nm and 60 nm. These shapes are consistent with VLP dimensions broadened laterally by imaging forces and tip curvature and reduced somewhat in height due to surface adhesion and possibly imaging forces. Although detailed characterization is subject of future work, we found that repeated imaging of VLPs with peak force of 1 nN led to gradual particle deformations: reduced particle heights and non-circular lateral cross-sections – consistent with VLP bursts (Figure 3C and Figure S6). This force is within the range of previously reported values (0.5-5 nN) for bursting various virus capsids^16, 17^ although most prior work has been done in liquid. Our observations represent an upper estimate of the rupture force for SARS-CoV-2. This demonstrates that SARS-CoV-2 VLPs can be disrupted by direct application of moderate mechanical perturbations and open the door for future studies of VLP mechanics and integrity.

**Figure 3.**
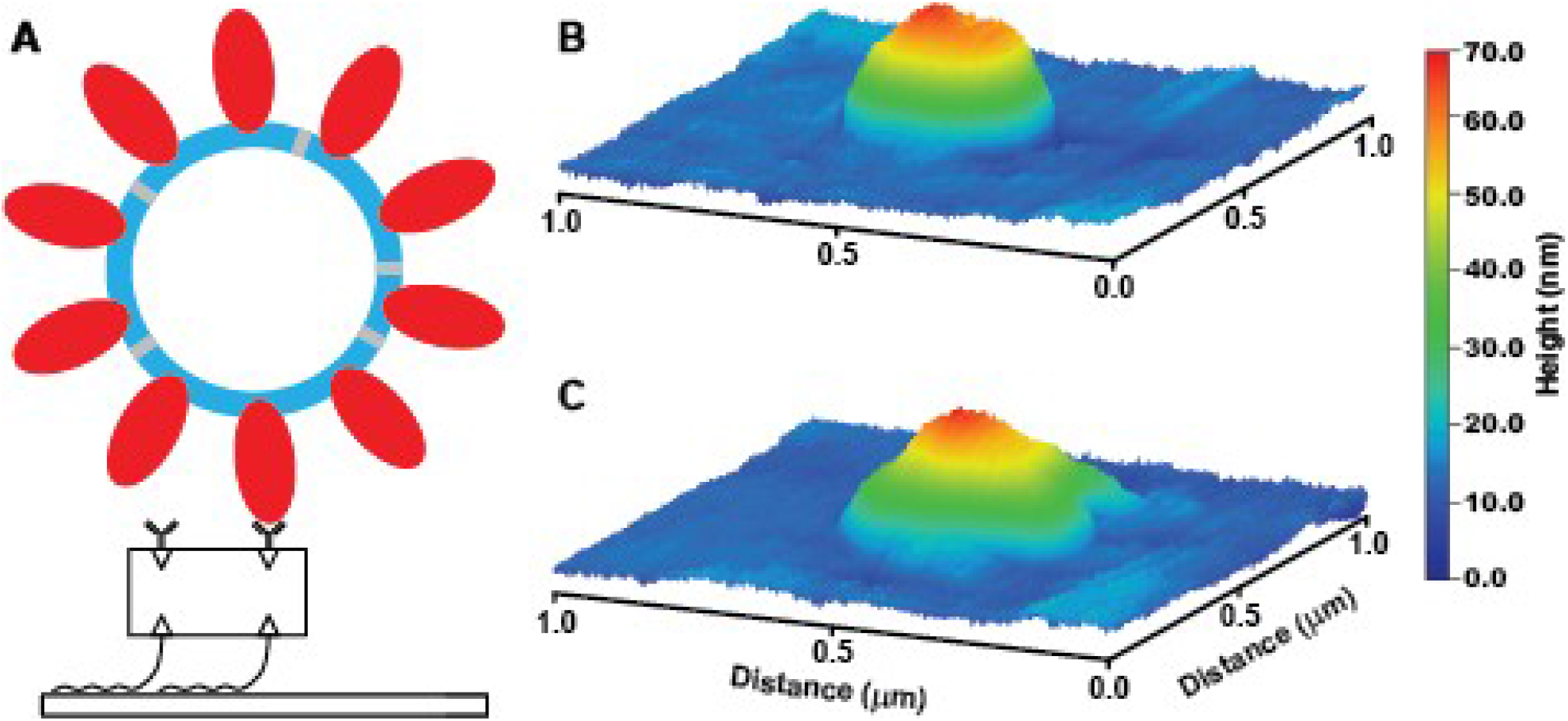
VLP surface immobilization and imaging. (a) Schematic of VLP-surface attachment strategy (bottom to top): glass surfaces were functionalized with Biotin-PEG-Silane, then neutravidin, then biotinylated anti-S antibody which then allowed for the capture of purified VLPs. VLP imaging showed initially symmetric particles (b) which developed prominent deformations likely indicative of bursts upon repeated AFM imaging with 1 nN peak force (c).

## Discussion

Expression of just M, S, and E proteins (Figure 1) was sufficient for release of VLPs which were structurally competent for both harvesting and subsequent investigations (Fig 2 and 3). These results open the door for many further studies. They can serve as immunogenic agents in place of the full infectious virus. The VLPs can also serve to study virus interactions with proteins of interest (e.g. receptors) – to date most such studies were only possible at the single protein level or in the context of the infectious virus. In addition, artificial VLPs can now be used to study the mechanical properties of the SARS-CoV-2 virus as well as their dependence on environmental conditions. Finally, VLPs can also be used to develop and validate novel testing strategies for SARS-CoV-2. The expression methodology is robust and should require no modifications to create VLPs for most S, M, and E mutations found in native conditions^18^.

## Availability

For expediency, genetic material in this work is published in the supplement and is available from the authors upon request. The authors will deposit the genetic material with Addgene.

## Acknowledgements

This study is supported by the NSF RAPID 2026657 award to MV and SS.

## Author contributions

HS performed the biochemical characterization AS performed the AFM characterization, BP, AP and CE created reagents, DB performed the electron microscopy characterization, MV and SS designed the research and wrote the manuscript.

## SUPPLEMENTARY INFORMATION

### Materials and Methods

#### Cell Culture

Cells from the human embryonic kidney 293T cell line were maintained in T-25 flasks using TrypLE Express Enzyme (Gibco) and DMEM with L-Glutamine, 4.5g/L glucose and sodium pyruvate (Corning) supplemented with 10% fetal bovine serum (Gibco). The cells were incubated at 37°C. Once over 90% confluent, the 293T cells were plated onto 100 mm tissue culture dishes with 10 mL of medium which would be ready for transfection the next day.

#### Transfection and VLP harvest

When the 293T cells were at 50% confluent in the 100 mm dishes, they were co-transfected with 29.5 μg of humanized membrane in pCDNA3.1, 5.9 μg of humanized spike in pCDNA3.1, and 5.9 μg of humanized envelope in pCDNA3.1 (GenScript). This transfection was carried out using 600 mL of Opti-MEM Reduced Serum Medium (Gibco) and 40 mL of Lipofectamine 2000 (Thermo Fisher Scientific). The cells were incubated at 37°C in 9 mL of Full DMEM with FBS supplementation for 24 hours.

#### Western Blot protocol

The specificity of the western blot antibodies are tested as shown in Figure S. the detailed protocol for western blot analysis is below:

1. Add sample buffer to the harvest. Add it in 1:1 ratio, i.e., 10ul sample+10ul buffer
2. Heat at 95°C for 5mins, while waiting set the gel.
3. After 5mins, quick spin the sample and start loading the gel
4. Run gel with running buffer at 90volt for 5mins and then at 160volt for 40mins or till the sample reaches the bottom of the gel
5. Prepare membrane and activate the PVDF membrane by methanol for 5mins. Put the foam and gel-blotting paper in transfect buffer for 5min. (2 foams and 4 blotting papers per gel)
6. Take out the gel and put it in transfer buffer
7. Wash the membrane with transfer buffer once before assembly
8. After you make the assembly stack run with transfer buffer at 35 volt (per gel) for 60mins
9. After the run, place the gel in a box with 7ml blocking buffer for 1hr
10. Add primary antibody with required dilution in 7 ml blocking buffer
11. Incubate overnight
12. Wash with PBS-Tween (2 times 15mins each)
13. Then add secondary antibody 1:20000. So, 0.25ul into 5ml of blocking buffer and keep for 45mins-1hr max and then again wash it with PBS-Tween (2 times 15mins each) and then with 15mins with PBS
14. Then visualize

#### Imaging

Licor Odyssey – I would need to double check the model if we need that information but I believe it is model 9120. You are correct that we image at 700 and 800 nm channels. We imaged at 0.0 mm focus offset. The image in Figure 1 was imaged at “highest quality” setting

**Figure S1.**
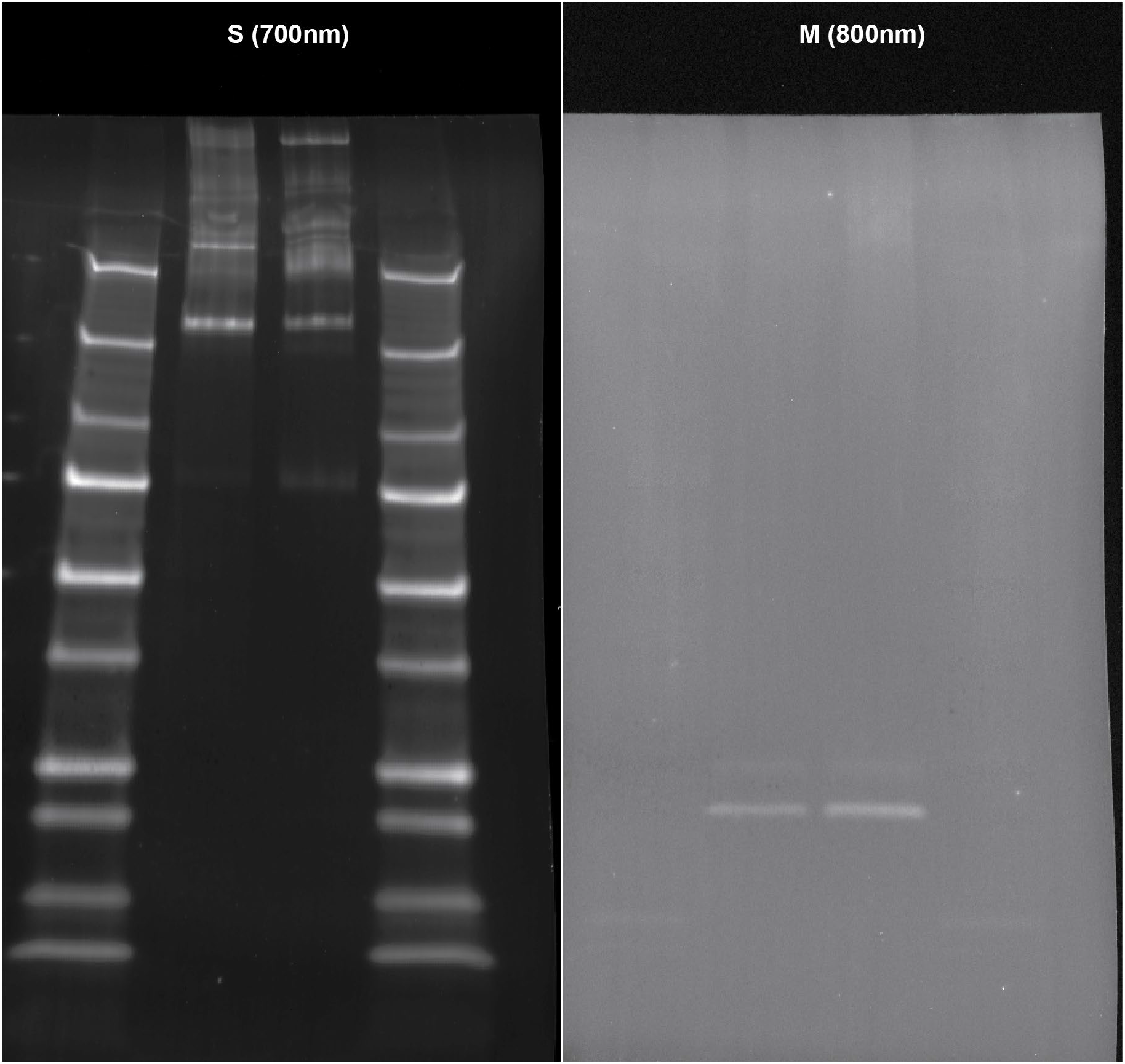
The data in Fig. 1B with channels separated. Here, no cropping or contrast adjustment was done. The original 16-bit TIF format images were resaved in 8-bit PNG format using ImageJ (NIH) without compression. Imaging channel labels were added for clarity.

**Figure S2.**
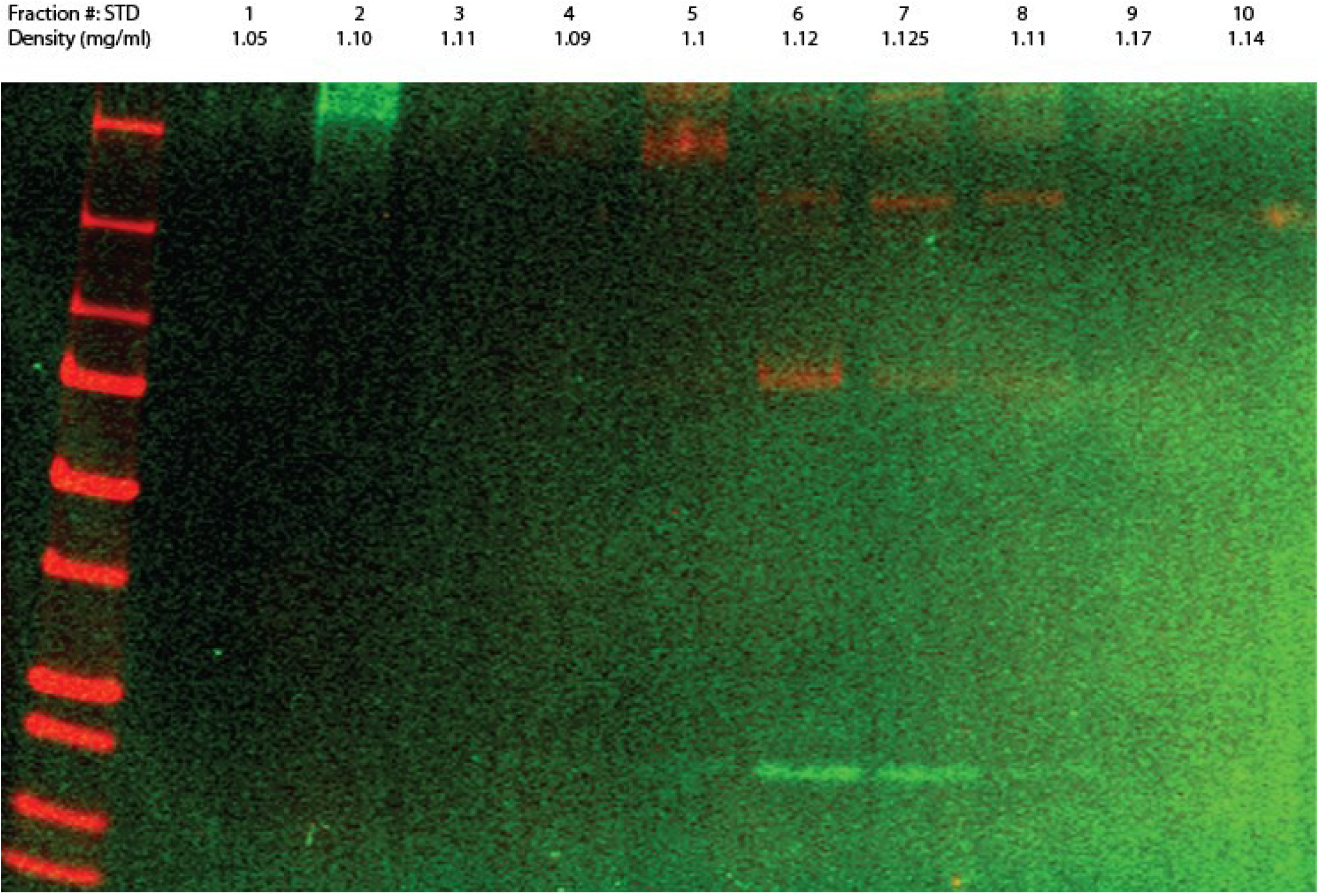
Sucrose fractionation of VLPs. The gel was performed simultaneously, scanned at 700 nm and 800 nm, then superimposed after minimal contrast enhancement. Protein size standards (leftmost lane) are same as Fig. 1B.

**Figure S3.**
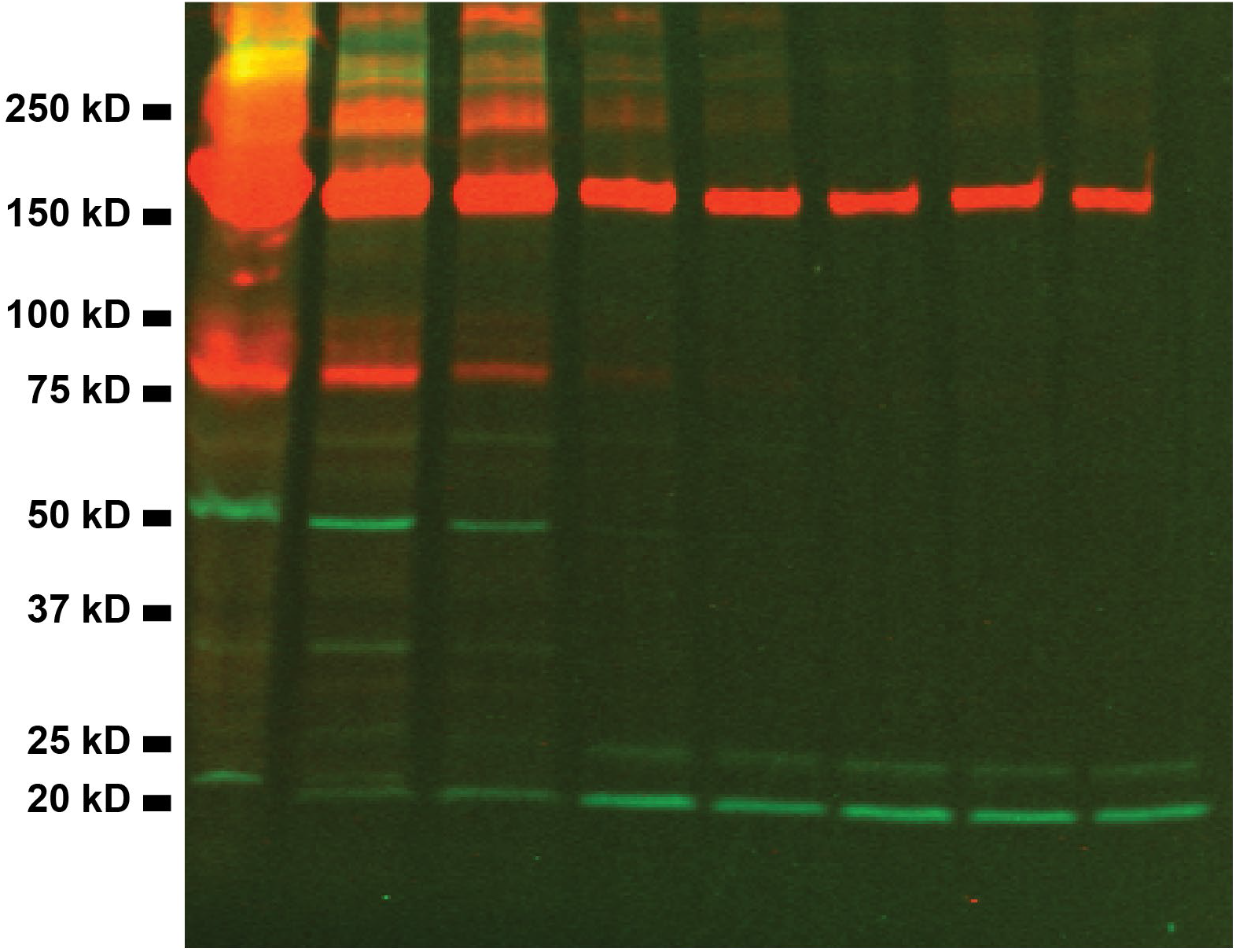
Serial 1:2 dilution analysis of cells. Both M (green) and S (red) proteins show molecular weight bands above their respective monomer bands (likely representative of aggregation) at highest concentrations. Monomer bands rapidly become dominant at lower dilutions.

**Figure S4.**
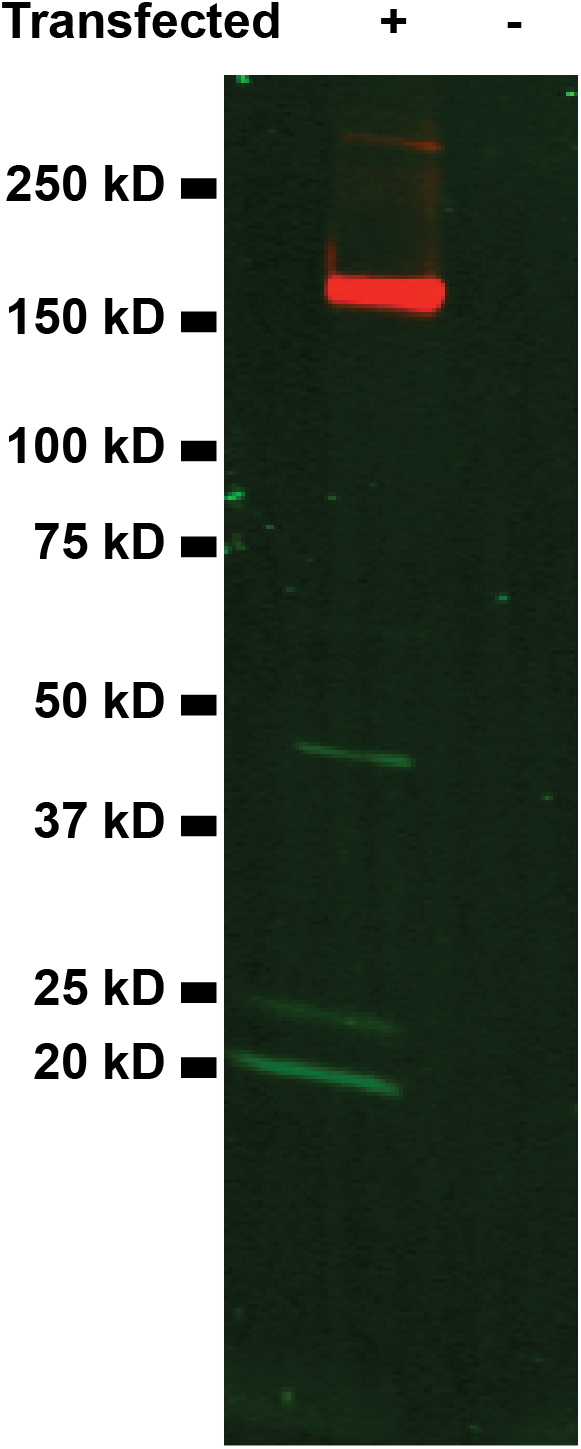
Anti-S and Anti-M antibodies are highly specific. Transfected cells show M and S presence via antibody probing however un-transfected cells show no bands under identical probing. Unrelated lanes are blanked out for clarity. The gel was performed simultaneously, scanned at 700 nm and 800 nm, then superimposed after minimal contrast enhancement. Protein size standards (leftmost lane) are same as Fig. 1B.

**Figure S5.**
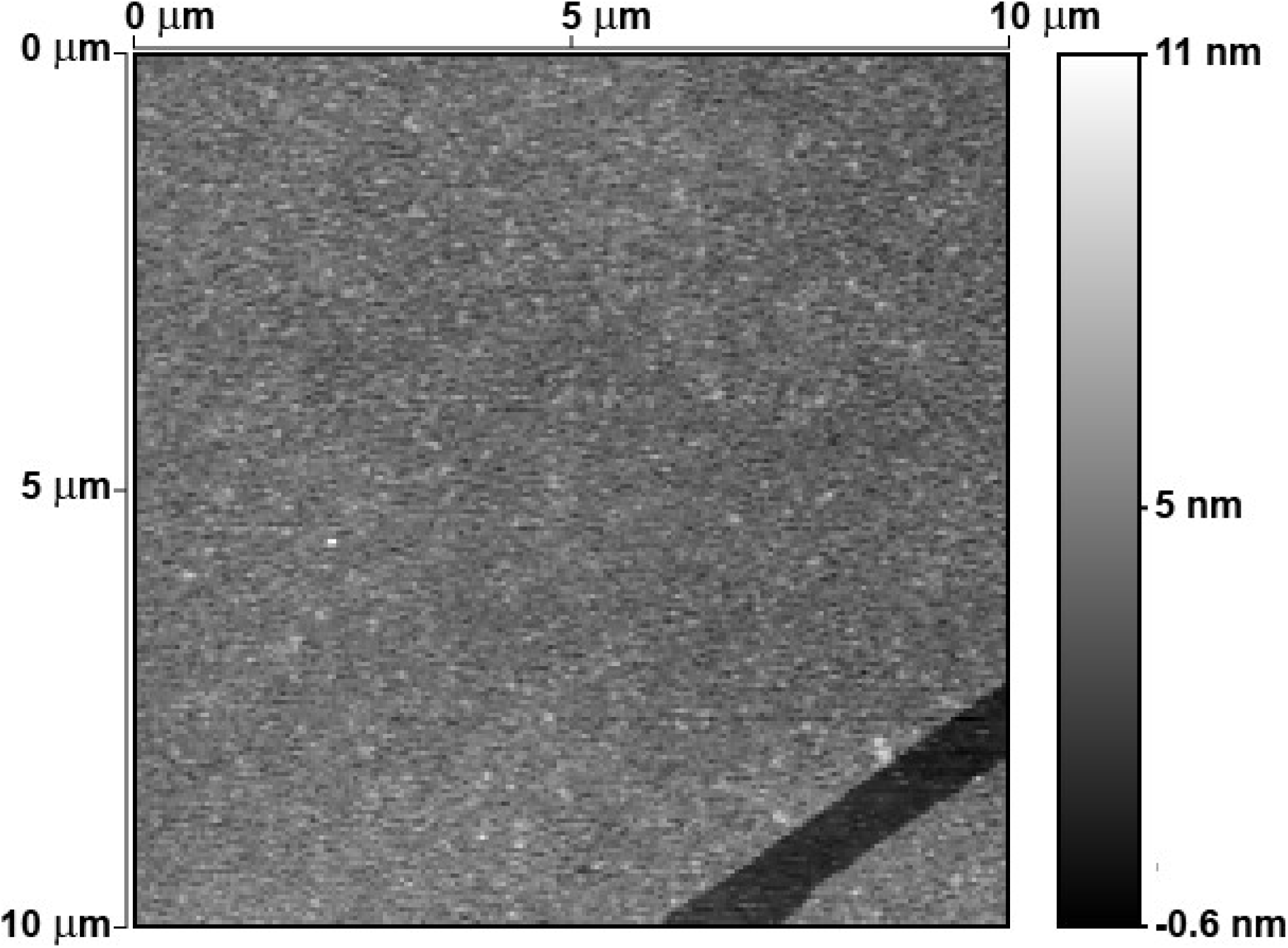
AFM imaging of glass surface functionalized with anti-S antibody. Glass surface functionalized as described in main text typically has large flat debris-free areas with occasional shallow step edges. These latter features are only seen after antibody adsorption and therefore correspond to submonolayer antibody coverage.

**Figure S6.**
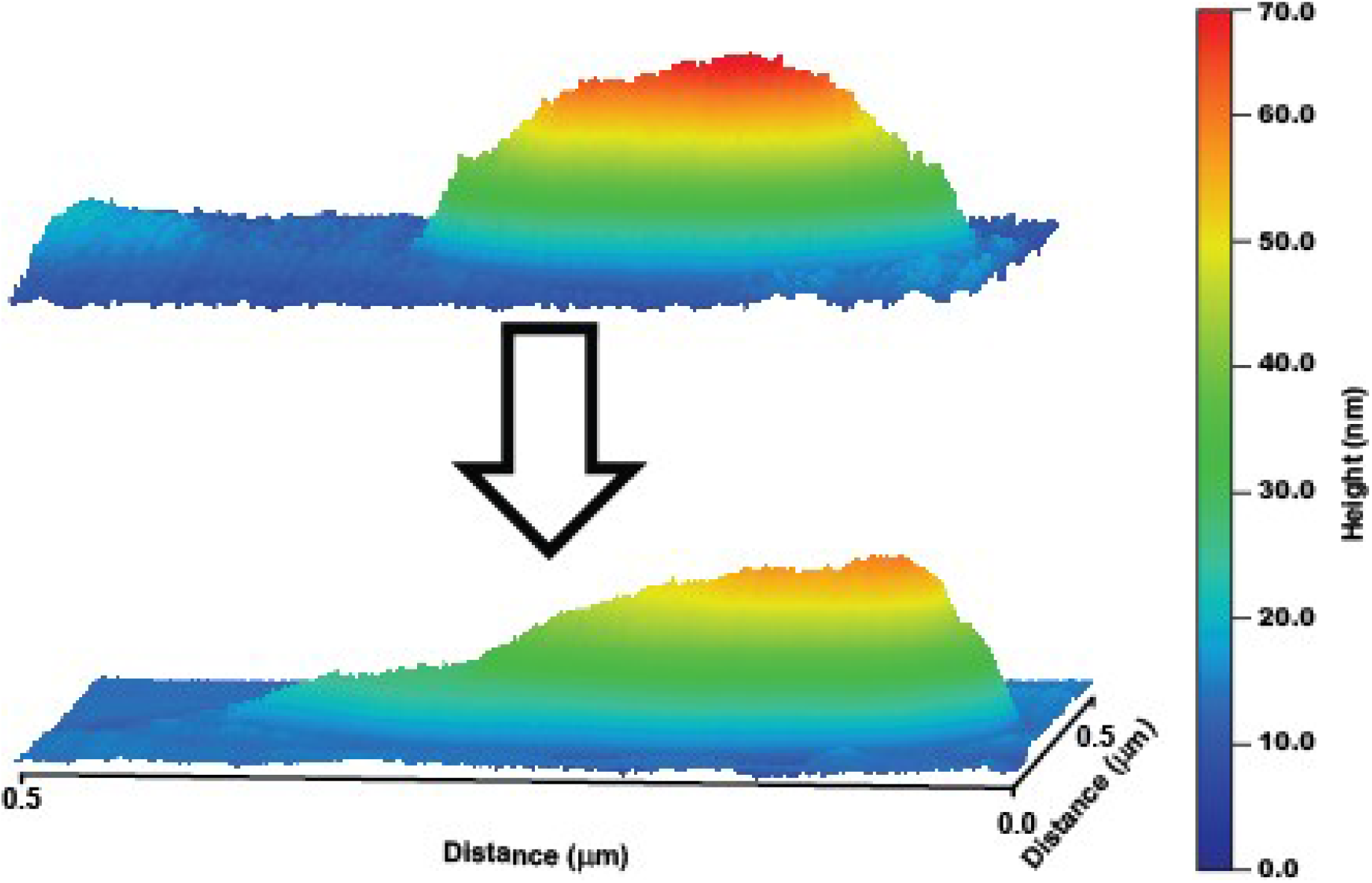
AFM imaging and bursting of a VLP. Several repeated scans at 1 nN peak force lead intact VLP (top) to burst (bottom).

#### Humanized S protein expression plasmid

GACGGATCGGGAGATCTCCCGATCCCCTATGGTGCACTCTCAGTACAATCTGCTCTGATGCCGCATAGTTAAGCCAGTATCTGCTCCCTGCTTGTGTGTTGGAGGTCGCTGAGTAGTGCGCGAGCAAAATTTAAGCTACAACAAGGCAAGGCTTGACCGACAATTGCATGAAGAATCTGCTTAGGGTTAGGCGTTTTGCGCTGCTTCGCGAGTACATTTATATTGGCTCATGTCCAATATGACCGCCATGTTGACATTGATTATTGACTAGTTATTAATAGTAATCAATTACGGGGTCATTAGTTCATAGCCCATATATGGAGTTCCGCGTTACATAACTTACGGTAAATGGCCCGCCTGGCTGACCGCCCAACGACCCCCGCCCATTGACGTCAATAATGACGTATGTTCCCATAGTAACGCCAATAGGGACTTTCCATTGACGTCAATGGGTGGAGTATTTACGGTAAACTGCCCACTTGGCAGTACATCAAGTGTATCATATGCCAAGTCCGCCCCCTATTGACGTCAATGACGGTAAATGGCCCGCCTGGCATTATGCCCAGTACATGACCTTACGGGACTTTCCTACTTGGCAGTACATCTACGTATTAGTCATCGCTATTACCATGGTGATGCGGTTTTGGCAGTACACCAATGGGCGTGGATAGCGGTTTGACTCACGGGGATTTCCAAGTCTCCACCCCATTGACGTCAATGGGAGTTTGTTTTGGCACCAAAATCAACGGGACTTTCCAAAATGTCGTAATAACCCCGCCCCGTTGACGCAAATGGGCGGTAGGCGTGTACGGTGGGAGGTCTATATAAGCAGAGCTCGTTTAGTGAACCGTCAGATCCTCACTCTCTTCCGCATCGCTGTCTGCGAGGGCCAGCTGTTGGGCTCGCGGTTGAGGACAAACTCTTCGCGGTCTTTCCAGTACTCTTGGATCGGAAACCCGTCGGCCTCCGAACGGTACTCCGCCACCGAGGGACCTGAGCGAGTCCGCATCGACCGGATCGGAAAACCTCTCGAGAAAGGCGTCTAACCAGTCACAGTCGCAAGGTAGGCTGAGCACCGTGGCGGGCGGCAGCGGGTGGCGGTCGGGGTTGTTTCTGGCGGAGGTGCTGCTGATGATGTAATTAAAGTAGGCGGTCTTGAGACGGCGGATGGTCGAGGTGAGGTGTGGCAGGCTTGAGATCCAGCTGTTGGGGTGAGTACTCCCTCTCAAAAGCGGGCATTACTTCTGCGCTAAGATTGTCAGTTTCCAAAAACGAGGAGGATTTGATATTCACCTGGCCCGATCTGGCCATACACTTGAGTGACAATGACATCCACTTTGCCTTTCTCTCCACAGGTGTCCACTCCCAGGTCCAAGTTTAAACTTTAATACGACTCACTATAGGGGCCGCCACCAAGCTTNNNNNGGTACCNNNNNATGTTTGTGTTCCTGGTGCTGCTGCCACTGGTGTCCAGCCAGTGTGTGAACCTGACCACCAGGACCCAACTTCCTCCTGCCTACACCAACTCCTTCACCAGGGGAGTCTACTACCCTGACAAGGTGTTCAGGTCCTCTGTGCTGCACAGCACCCAGGACCTGTTCCTGCCATTCTTCAGCAATGTGACCTGGTTCCATGCCATCCATGTGTCTGGCACCAATGGCACCAAGAGGTTTGACAACCCTGTGCTGCCATTCAATGATGGAGTCTACTTTGCCAGCACAGAGAAGAGCAACATCATCAGGGGCTGGATTTTTGGCACCACCCTGGACAGCAAGACCCAGTCCCTGCTGATTGTGAACAATGCCACCAATGTGGTGATTAAGGTGTGTGAGTTCCAGTTCTGTAATGACCCATTCCTGGGAGTCTACTACCACAAGAACAACAAGTCCTGGATGGAGTCTGAGTTCAGGGTCTACTCCTCTGCCAACAACTGTACCTTTGAATATGTGAGCCAACCATTCCTGATGGACTTGGAGGGCAAGCAGGGCAACTTCAAGAACCTGAGGGAGTTTGTGTTCAAGAACATTGATGGCTACTTCAAGATTTACAGCAAACACACACCAATCAACCTGGTGAGGGACCTGCCACAGGGCTTCTCTGCCTTGGAACCACTGGTGGACCTGCCAATTGGCATCAACATCACCAGGTTCCAGACCCTGCTGGCTCTGCACAGGTCCTACCTGACACCTGGAGACTCCTCCTCTGGCTGGACAGCAGGAGCAGCAGCCTACTATGTGGGCTACCTCCAACCAAGGACCTTCCTGCTGAAATACAATGAGAATGGCACCATCACAGATGCTGTGGACTGTGCCCTGGACCCACTGTCTGAGACCAAGTGTACCCTGAAATCCTTCACAGTGGAGAAGGGCATCTACCAGACCAGCAACTTCAGGGTCCAACCAACAGAGAGCATTGTGAGGTTTCCAAACATCACCAACCTGTGTCCATTTGGAGAGGTGTTCAATGCCACCAGGTTTGCCTCTGTCTATGCCTGGAACAGGAAGAGGATTAGCAACTGTGTGGCTGACTACTCTGTGCTCTACAACTCTGCCTCCTTCAGCACCTTCAAGTGTTATGGAGTGAGCCCAACCAAACTGAATGACCTGTGTTTCACCAATGTCTATGCTGACTCCTTTGTGATTAGGGGAGATGAGGTGAGACAGATTGCCCCTGGACAAACAGGCAAGATTGCTGACTACAACTACAAACTGCCTGATGACTTCACAGGCTGTGTGATTGCCTGGAACAGCAACAACCTGGACAGCAAGGTGGGAGGCAACTACAACTACCTCTACAGACTGTTCAGGAAGAGCAACCTGAAACCATTTGAGAGGGACATCAGCACAGAGATTTACCAGGCTGGCAGCACACCATGTAATGGAGTGGA GGGCTTCAACTGTTACTTTCCACTCCAATCCTATGGCTTCCAACCAACCAATGGAGTGGGCTACCAACCATACAGGGTGGTGGTGCTGTCCTTTGAACTGCTCCATGCCCCTGCCACAGTGTGTGGACCAAAGAAGAGCACCAACCTGGTGAAGAACAAGTGTGTGAACTTCAACTTCAATGGACTGACAGGCACAGGAGTGCTGACAGAGAGCAACAAGAAGTTCCTGCCATTCCAACAGTTTGGCAGGGACATTGCTGACACCACAGATGCTGTGAGGGACCCACAGACCTTGGAGATTCTGGACATCACACCATGTTCCTTTGGAGGAGTGTCTGTGATTACACCTGGCACCAACACCAGCAACCAGGTGGCTGTGCTCTACCAGGATGTGAACTGTACTGAGGTGCCTGTGGCTATCCATGCTGACCAACTTACACCAACCTGGAGGGTCTACAGCACAGGCAGCAATGTGTTCCAGACCAGGGCTGGCTGTCTGATTGGAGCAGAGCATGTGAACAACTCCTATGAGTGTGACATCCCAATTGGAGCAGGCATCTGTGCCTCCTACCAGACCCAGACCAACAGCCCAAGGAGGGCAAGGTCTGTGGCAAGCCAGAGCATCATTGCCTACACAATGAGTCTGGGAGCAGAGAACTCTGTGGCTTACAGCAACAACAGCATTGCCATCCCAACCAACTTCACCATCTCTGTGACCACAGAGATTCTGCCTGTGAGTATGACCAAGACCTCTGTGGACTGTACAATGTATATCTGTGGAGACAGCACAGAGTGTAGCAACCTGCTGCTCCAATATGGCTCCTTCTGTACCCAACTTAACAGGGCTCTGACAGGCATTGCTGTGGAACAGGACAAGAACACCCAGGAGGTGTTTGCCCAGGTGAAGCAGATTTACAAGACACCTCCAATCAAGGACTTTGGAGGCTTCAACTTCAGCCAGATTCTGCCTGACCCAAGCAAGCCAAGCAAGAGGTCCTTCATTGAGGACCTGCTGTTCAACAAGGTGACCCTGGCTGATGCTGGCTTCATCAAGCAATATGGAGACTGTCTGGGAGACATTGCTGCCAGGGACCTGATTTGTGCCCAGAAGTTCAATGGACTGACAGTGCTGCCTCCACTGCTGACAGATGAGATGATTGCCCAATACACCTCTGCCCTGCTGGCTGGCACCATCACCTCTGGCTGGACCTTTGGAGCAGGAGCAGCCCTCCAAATCCCATTTGCTATGCAGATGGCTTACAGGTTCAATGGCATTGGAGTGACCCAGAATGTGCTCTATGAGAACCAGAAACTGATTGCCAACCAGTTCAACTCTGCCATTGGCAAGATTCAGGACTCCCTGTCCAGCACAGCCTCTGCCCTGGGCAAACTCCAAGATGTGGTGAACCAGAATGCCCAGGCTCTGAACACCCTGGTGAAGCAACTTTCCAGCAACTTTGGAGCCATCTCCTCTGTGCTGAATGACATCCTGAGCAGACTGGACAAGGTGGAGGCTGAGGTCCAGATTGACAGACTGATTACAGGCAGACTCCAATCCCTCCAAACCTATGTGACCCAACAACTTATCAGGGCTGCTGAGATTAGGGCATCTGCCAACCTGGCTGCCACCAAGATGAGTGAGTGTGTGCTGGGACAAAGCAAGAGGGTGGACTTCTGTGGCAAGGGCTACCACCTGATGAGTTTTCCACAGTCTGCCCCTCATGGAGTGGTGTTCCTGCATGTGACCTATGTGCCTGCCCAGGAGAAGAACTTCACCACAGCCCCTGCCATCTGCCATGATGGCAAGGCTCACTTTCCAAGGGAGGGAGTGTTTGTGAGCAATGGCACCCACTGGTTTGTGACCCAGAGGAACTTCTATGAACCACAGATTATCACCACAGACAACACCTTTGTGTCTGGCAACTGTGATGTGGTGATTGGCATTGTGAACAACACAGTCTATGACCCACTCCAACCTGAACTGGACTCCTTCAAGGAGGAACTGGACAAATACTTCAAGAACCACACCAGCCCTGATGTGGACCTGGGAGACATCTCTGGCATCAATGCCTCTGTGGTGAACATCCAGAAGGAGATTGACAGACTGAATGAGGTGGCTAAGAACCTGAATGAGTCCCTGATTGACCTCCAAGAACTGGGCAAATATGAACAATACATCAAGTGGCCATGGTACATCTGGCTGGGCTTCATTGCTGGACTGATTGCCATTGTGATGGTGACCATAATGCTGTGTTGTATGACCTCCTGTTGTTCCTGTCTGAAAGGCTGTTGTTCCTGTGGCTCCTGTTGTAAGTTTGATGAGGATGACTCTGAACCTGTGCTGAAAGGAGTGAAACTGCACTACACCTGANNNNNTCTAGANNNNNGCGGCCGCCGAATTCGGGCCCGTTTAAACCCGCTGATCAGCCTCGACTGTGCCTTCTAGTTGCCAGCCATCTGTTGTTTGCCCCTCCCCCGTGCCTTCCTTGACCCTGGAAGGTGCCACTCCCACTGTCCTTTCCTAATAAAATGAGGAAATTGCATCGCATTGTCTGAGTAGGTGTCATTCTATTCTGGGGGGTGGGGTGGGGCAGGACAGCAAGGGGGAGGATTGGGAAGACAATAGCAGGCATGCTGGGGATGCGGTGGGCTCTATGGCTTCTGAGGCGGAAAGAACCAGCTGGGGCTCTAGGGGGTATCCCCACGCGCCCTGTAGCGGCGCATTAAGCGCGGCGGGTGTGGTGGTTACGCGCAGCGTGACCGCTACACTTGCCAGCGCCCTAGCGCCCGCTCCTTTCGCTTTCTTCCCTTCCTTTCTCGCCACGTTCGCCGGCTTTCCCCGTCAAGCTCTAAATCGGGGGCTCCCTTTAGGGTTCCGATTTAGTGCTTTACGGCACCTCGACCCCAAAAAACTTGATTAGGGTGATGGTTCACGTAGTGGGCCATCGCCCTGATAGACGGTTTT TCGCCCTTTGACGTTGGAGTCCACGTTCTTTAATAGTGGACTCTTGTTCCAAACTGGAACAACACTCAACCCTATCTCGGTCTATTCTTTTGATTTATAAGGGATTTTGCCGATTTCGGCCTATTGGTTAAAAAATGAGCTGATTTAACAAAAATTTAACGCGAATTAATTCTGTGGAATGTGTGTCAGTTAGGGTGTGGAAAGTCCCCAGGCTCCCCAGCAGGCAGAAGTATGCAAAGCATGCATCTCAATTAGTCAGCAACCAGGTGTGGAAAGTCCCCAGGCTCCCCAGCAGGCAGAAGTATGCAAAGCATGCATCTCAATTAGTCAGCAACCATAGTCCCGCCCCTAACTCCGCCCATCCCGCCCCTAACTCCGCCCAGTTCCGCCCATTCTCCGCCCCATGGCTGACTAATTTTTTTTATTTATGCAGAGGCCGAGGCCGCCTCTGCCTCTGAGCTATTCCAGAAGTAGTGAGGAGGCTTTTTTGGAGGCCTAGGCTTTTGCAAAAAGCTCCCGGGAGCTTGTATATCCATTTTCGGATCTGATCAGCACGTGATGAAAAAGCCTGAACTCACCGCGACGTCTGTCGAGAAGTTTCTGATCGAAAAGTTCGACAGCGTCTCCGACCTGATGCAGCTCTCGGAGGGCGAAGAATCTCGTGCTTTCAGCTTCGATGTAGGAGGGCGTGGATATGTCCTGCGGGTAAATAGCTGCGCCGATGGTTTCTACAAAGATCGTTATGTTTATCGGCACTTTGCATCGGCCGCGCTCCCGATTCCGGAAGTGCTTGACATTGGGGAATTCAGCGAGAGCCTGACCTATTGCATCTCCCGCCGTGCACAGGGTGTCACGTTGCAAGACCTGCCTGAAACCGAACTGCCCGCTGTTCTGCAGCCGGTCGCGGAGGCCATGGATGCGATCGCTGCGGCCGATCTTAGCCAGACGAGCGGGTTCGGCCCATTCGGACCGCAAGGAATCGGTCAATACACTACATGGCGTGATTTCATATGCGCGATTGCTGATCCCCATGTGTATCACTGGCAAACTGTGATGGACGACACCGTCAGTGCGTCCGTCGCGCAGGCTCTCGATGAGCTGATGCTTTGGGCCGAGGACTGCCCCGAAGTCCGGCACCTCGTGCACGCGGATTTCGGCTCCAACAATGTCCTGACGGACAATGGCCGCATAACAGCGGTCATTGACTGGAGCGAGGCGATGTTCGGGGATTCCCAATACGAGGTCGCCAACATCTTCTTCTGGAGGCCGTGGTTGGCTTGTATGGAGCAGCAGACGCGCTACTTCGAGCGGAGGCATCCGGAGCTTGCAGGATCGCCGCGGCTCCGGGCGTATATGCTCCGCATTGGTCTTGACCAACTCTATCAGAGCTTGGTTGACGGCAATTTCGATGATGCAGCTTGGGCGCAGGGTCGATGCGACGCAATCGTCCGATCCGGAGCCGGGACTGTCGGGCGTACACAAATCGCCCGCAGAAGCGCGGCCGTCTGGACCGATGGCTGTGTAGAAGTACTCGCCGATAGTGGAAACCGACGCCCCAGCACTCGTCCGAGGGCAAAGGAATAGCACGTGCTACGAGATTTCGATTCCACCGCCGCCTTCTATGAAAGGTTGGGCTTCGGAATCGTTTTCCGGGACGCCGGCTGGATGATCCTCCAGCGCGGGGATCTCATGCTGGAGTTCTTCGCCCACCCCAACTTGTTTATTGCAGCTTATAATGGTTACAAATAAAGCAATAGCATCACAAATTTCACAAATAAAGCATTTTTTTCACTGCATTCTAGTTGTGGTTTGTCCAAACTCATCAATGTATCTTATCATGTCTGTATACCGTCGACCTCTAGCTAGAGCTTGGCGTAATCATGGTCATAGCTGTTTCCTGTGTGAAATTGTTATCCGCTCACAATTCCACACAACATACGAGCCGGAAGCATAAAGTGTAAAGCCTGGGGTGCCTAATGAGTGAGCTAACTCACATTAATTGCGTTGCGCTCACTGCCCGCTTTCCAGTCGGGAAACCTGTCGTGCCAGCTGCATTAATGAATCGGCCAACGCGCGGGGAGAGGCGGTTTGCGTATTGGGCGCTCTTCCGCTTCCTCGCTCACTGACTCGCTGCGCTCGGTCGTTCGGCTGCGGCGAGCGGTATCAGCTCACTCAAAGGCGGTAATACGGTTATCCACAGAATCAGGGGATAACGCAGGAAAGAACATGTGAGCAAAAGGCCAGCAAAAGGCCAGGAACCGTAAAAAGGCCGCGTTGCTGGCGTTTTTCCATAGGCTCCGCCCCCCTGACGAGCATCACAAAAATCGACGCTCAAGTCAGAGGTGGCGAAACCCGACAGGACTATAAAGATACCAGGCGTTTCCCCCTGGAAGCTCCCTCGTGCGCTCTCCTGTTCCGACCCTGCCGCTTACCGGATACCTGTCCGCCTTTCTCCCTTCGGGAAGCGTGGCGCTTTCTCATAGCTCACGCTGTAGGTATCTCAGTTCGGTGTAGGTCGTTCGCTCCAAGCTGGGCTGTGTGCACGAACCCCCCGTTCAGCCCGACCGCTGCGCCTTATCCGGTAACTATCGTCTTGAGTCCAACCCGGTAAGACACGACTTATCGCCACTGGCAGCAGCCACTGGTAACAGGATTAGCAGAGCGAGGTATGTAGGCGGTGCTACAGAGTTCTTGAAGTGGTGGCCTAACTACGGCTACACTAGAAGAACAGTATTTGGTATCTGCGCTCTGCTGAAGCCAGTTACCTTCGGAAAAAGAGTTGGTAGCTCTTGATCCGGCAAACAAACCACCGCTGGTAGCGGTTTTTTTGTTTGCAAGCAGCAGATTACGCGCAGAAAAAAAGGATCTCAAGAAGATCCTTTGATCTTTTCTACGGGGTCTGACGCTCAGTGGAACGAAAACTCACGTTAAGGGATTTTGGTCATGAGATTATCAAAAAGGATCTTCACCTA GATCCTTTTAAATTAAAAATGAAGTTTTAAATCAATCTAAAGTATATATGAGTAAACTTGGTCTGACAGTTACCAATGCTTAATCAGTGAGGCACCTATCTCAGCGATCTGTCTATTTCGTTCATCCATAGTTGCCTGACTCCCCGTCGTGTAGATAACTACGATACGGGAGGGCTTACCATCTGGCCCCAGTGCTGCAATGATACCGCGAGACCCACGCTCACCGGCTCCAGATTTATCAGCAATAAACCAGCCAGCCGGAAGGGCCGAGCGCAGAAGTGGTCCTGCAACTTTATCCGCCTCCATCCAGTCTATTAATTGTTGCCGGGAAGCTAGAGTAAGTAGTTCGCCAGTTAATAGTTTGCGCAACGTTGTTGCCATTGCTACAGGCATCGTGGTGTCACGCTCGTCGTTTGGTATGGCTTCATTCAGCTCCGGTTCCCAACGATCAAGGCGAGTTACATGATCCCCCATGTTGTGCAAAAAAGCGGTTAGCTCCTTCGGTCCTCCGATCGTTGTCAGAAGTAAGTTGGCCGCAGTGTTATCACTCATGGTTATGGCAGCACTGCATAATTCTCTTACTGTCATGCCATCCGTAAGATGCTTTTCTGTGACTGGTGAGTACTCAACCAAGTCATTCTGAGAATAGTGTATGCGGCGACCGAGTTGCTCTTGCCCGGCGTCAATACGGGATAATACCGCGCCACATAGCAGAACTTTAAAAGTGCTCATCATTGGAAAACGTTCTTCGGGGCGAAAACTCTCAAGGATCTTACCGCTGTTGAGATCCAGTTCGATGTAACCCACTCGTGCACCCAACTGATCTTCAGCATCTTTTACTTTCACCAGCGTTTCTGGGTGAGCAAAAACAGGAAGGCAAAATGCCGCAAAAAAGGGAATAAGGGCGACACGGAAATGTTGAATACTCATACTCTTCCTTTTTCAATATTATTGAAGCATTTATCAGGGTTATTGTCTCATGAGCGGATACATATTTGAATGTATTTAGAAAAATAAACAAATAGGGGTTCCGCGCACATTTCCCCGAAAAGTGCCACCTGACGTC

#### Humanized M protein expression plasmid

GACGGATCGGGAGATCTCCCGATCCCCTATGGTGCACTCTCAGTACAATCTGCTCTGATGCCGCATAGTTAAGCCAGTATCTGCTCCCTGCTTGTGTGTTGGAGGTCGCTGAGTAGTGCGCGAGCAAAATTTAAGCTACAACAAGGCAAGGCTTGACCGACAATTGCATGAAGAATCTGCTTAGGGTTAGGCGTTTTGCGCTGCTTCGCGATGTACGGGCCAGATATACGCGTTGACATTGATTATTGACTAGTTATTAATAGTAATCAATTACGGGGTCATTAGTTCATAGCCCATATATGGAGTTCCGCGTTACATAACTTACGGTAAATGGCCCGCCTGGCTGACCGCCCAACGACCCCCGCCCATTGACGTCAATAATGACGTATGTTCCCATAGTAACGCCAATAGGGACTTTCCATTGACGTCAATGGGTGGAGTATTTACGGTAAACTGCCCACTTGGCAGTACATCAAGTGTATCATATGCCAAGTACGCCCCCTATTGACGTCAATGACGGTAAATGGCCCGCCTGGCATTATGCCCAGTACATGACCTTATGGGACTTTCCTACTTGGCAGTACATCTACGTATTAGTCATCGCTATTACCATGGTGATGCGGTTTTGGCAGTACATCAATGGGCGTGGATAGCGGTTTGACTCACGGGGATTTCCAAGTCTCCACCCCATTGACGTCAATGGGAGTTTGTTTTGGCACCAAAATCAACGGGACTTTCCAAAATGTCGTAACAACTCCGCCCCATTGACGCAAATGGGCGGTAGGCGTGTACGGTGGGAGGTCTATATAAGCAGAGCTCTCTGGCTAACTAGAGAACCCACTGCTTACTGGCTTATCGAAATTAATACGACTCACTATAGGGAGACCCAAGCTGGCTAGCGCAGCATCTTCAGCTCTTGGGTTTAGCAAAAGTTCATCTCCTTCTGCATCCTTAACTGAGAATGAGCTTTTGTGGGAGCCCACACCAGTCAAGTTGGATTTGAACCCAGCTGCTCTGTACAAGCTTGGTACCGCCACCATGGCCGACAGCAACGGCACCATCACCGTGGAGGAGCTGAAGAAGCTGCTGGAGCAGTGGAACCTGGTGATCGGCTTCCTGTTCCTGACCTGGATCTGCCTGCTGCAGTTCGCCTACGCCAACAGGAACAGGTTCCTGTACATCATCAAGCTGATCTTCCTGTGGCTGCTGTGGCCCGTGACCCTGGCCTGCTTCGTGCTGGCCGCCGTGTACAGGATCAACTGGATCACCGGCGGCATCGCCATCGCCATGGCCTGCCTGGTGGGCCTGATGTGGCTGAGCTACTTCATCGCCAGCTTCAGGCTGTTCGCCAGGACCAGGAGCATGTGGAGCTTCAACCCCGAGACCAACATCCTGCTGAACGTGCCCCTGCACGGCACCATCCTGACCAGGCCCCTGCTGGAGAGCGAGCTGGTGATCGGCGCCGTGATCCTGAGGGGCCACCTGAGGATCGCCGGCCACCACCTGGGCAGGTGCGACATCAAGGACCTGCCCAAGGAGATCACCGTGGCCACCAGCAGGACCCTGAGCTACTACAAGCTGGGCGCCAGCCAGAGGGTGGCCGGCGACAGCGGCTTCGCCGCCTACAGCAGGTACAGGATCGGCAACTACAAGCTGAA CACCGACCACAGCAGCAGCAGCGACAACATCGCCCTGCTGGTGCAGTAATCTAGAGGGCCCGTTTAAACCCGCTGATCAGCCTCGACTGTGCCTTCTAGTTGCCAGCCATCTGTTGTTTGCCCCTCCCCCGTGCCTTCCTTGACCCTGGAAGGTGCCACTCCCACTGTCCTTTCCTAATAAAATGAGGAAATTGCATCGCATTGTCTGAGTAGGTGTCATTCTATTCTGGGGGGTGGGGTGGGGCAGGACAGCAAGGGGGAGGATTGGGAAGACAATAGCAGGCATGCTGGGGATGCGGTGGGCTCTATGGCTTCTGAGGCGGAAAGAACCAGCTGGGGCTCTAGGGGGTATCCCCACGCGCCCTGTAGCGGCGCATTAAGCGCGGCGGGTGTGGTGGTTACGCGCAGCGTGACCGCTACACTTGCCAGCGCCCTAGCGCCCGCTCCTTTCGCTTTCTTCCCTTCCTTTCTCGCCACGTTCGCCGGCTTTCCCCGTCAAGCTCTAAATCGGGGGCTCCCTTTAGGGTTCCGATTTAGTGCTTTACGGCACCTCGACCCCAAAAAACTTGATTAGGGTGATGGTTCACGTAGTGGGCCATCGCCCTGATAGACGGTTTTTCGCCCTTTGACGTTGGAGTCCACGTTCTTTAATAGTGGACTCTTGTTCCAAACTGGAACAACACTCAACCCTATCTCGGTCTATTCTTTTGATTTATAAGGGATTTTGCCGATTTCGGCCTATTGGTTAAAAAATGAGCTGATTTAACAAAAATTTAACGCGAATTAATTCTGTGGAATGTGTGTCAGTTAGGGTGTGGAAAGTCCCCAGGCTCCCCAGCAGGCAGAAGTATGCAAAGCATGCATCTCAATTAGTCAGCAACCAGGTGTGGAAAGTCCCCAGGCTCCCCAGCAGGCAGAAGTATGCAAAGCATGCATCTCAATTAGTCAGCAACCATAGTCCCGCCCCTAACTCCGCCCATCCCGCCCCTAACTCCGCCCAGTTCCGCCCATTCTCCGCCCCATGGCTGACTAATTTTTTTTATTTATGCAGAGGCCGAGGCCGCCTCTGCCTCTGAGCTATTCCAGAAGTAGTGAGGAGGCTTTTTTGGAGGCCTAGGCTTTTGCAAAAAGCTCCCGGGAGCTTGTATATCCATTTTCGGATCTGATCAAGAGACAGGATGAGGATCGTTTCGCATGATTGAACAAGATGGATTGCACGCAGGTTCTCCGGCCGCTTGGGTGGAGAGGCTATTCGGCTATGACTGGGCACAACAGACAATCGGCTGCTCTGATGCCGCCGTGTTCCGGCTGTCAGCGCAGGGGCGCCCGGTTCTTTTTGTCAAGACCGACCTGTCCGGTGCCCTGAATGAACTGCAGGACGAGGCAGCGCGGCTATCGTGGCTGGCCACGACGGGCGTTCCTTGCGCAGCTGTGCTCGACGTTGTCACTGAAGCGGGAAGGGACTGGCTGCTATTGGGCGAAGTGCCGGGGCAGGATCTCCTGTCATCTCACCTTGCTCCTGCCGAGAAAGTATCCATCATGGCTGATGCAATGCGGCGGCTGCATACGCTTGATCCGGCTACCTGCCCATTCGACCACCAAGCGAAACATCGCATCGAGCGAGCACGTACTCGGATGGAAGCCGGTCTTGTCGATCAGGATGATCTGGACGAAGAGCATCAGGGGCTCGCGCCAGCCGAACTGTTCGCCAGGCTCAAGGCGCGCATGCCCGACGGCGAGGATCTCGTCGTGACCCATGGCGATGCCTGCTTGCCGAATATCATGGTGGAAAATGGCCGCTTTTCTGGATTCATCGACTGTGGCCGGCTGGGTGTGGCGGACCGCTATCAGGACATAGCGTTGGCTACCCGTGATATTGCTGAAGAGCTTGGCGGCGAATGGGCTGACCGCTTCCTCGTGCTTTACGGTATCGCCGCTCCCGATTCGCAGCGCATCGCCTTCTATCGCCTTCTTGACGAGTTCTTCTGAGCGGGACTCTGGGGTTCGAAATGACCGACCAAGCGACGCCCAACCTGCCATCACGAGATTTCGATTCCACCGCCGCCTTCTATGAAAGGTTGGGCTTCGGAATCGTTTTCCGGGACGCCGGCTGGATGATCCTCCAGCGCGGGGATCTCATGCTGGAGTTCTTCGCCCACCCCAACTTGTTTATTGCAGCTTATAATGGTTACAAATAAAGCAATAGCATCACAAATTTCACAAATAAAGCATTTTTTTCACTGCATTCTAGTTGTGGTTTGTCCAAACTCATCAATGTATCTTATCATGTCTGTATACCGTCGACCTCTAGCTAGAGCTTGGCGTAATCATGGTCATAGCTGTTTCCTGTGTGAAATTGTTATCCGCTCACAATTCCACACAACATACGAGCCGGAAGCATAAAGTGTAAAGCCTGGGGTGCCTAATGAGTGAGCTAACTCACATTAATTGCGTTGCGCTCACTGCCCGCTTTCCAGTCGGGAAACCTGTCGTGCCAGCTGCATTAATGAATCGGCCAACGCGCGGGGAGAGGCGGTTTGCGTATTGGGCGCTCTTCCGCTTCCTCGCTCACTGACTCGCTGCGCTCGGTCGTTCGGCTGCGGCGAGCGGTATCAGCTCACTCAAAGGCGGTAATACGGTTATCCACAGAATCAGGGGATAACGCAGGAAAGAACATGTGAGCAAAAGGCCAGCAAAAGGCCAGGAACCGTAAAAAGGCCGCGTTGCTGGCGTTTTTCCATAGGCTCCGCCCCCCTGACGAGCATCACAAAAATCGACGCTCAAGTCAGAGGTGGCGAAACCCGACAGGACTATAAAGATACCAGGCGTTTCCCCCTGGAAGCTCCCTCGTGCGCTCTCCTGTTCCGACCCTGCCGCTTACCGGATACCTGTCCGCCTTTCTCCCTTCGGGAAGCGTGGCGCTTTCTCATAGCTCACGCTGTAGGTATCTCAGTTCGGTGTAGGTCGTTCGCT CCAAGCTGGGCTGTGTGCACGAACCCCCCGTTCAGCCCGACCGCTGCGCCTTATCCGGTAACTATCGTCTTGAGTCCAACCCGGTAAGACACGACTTATCGCCACTGGCAGCAGCCACTGGTAACAGGATTAGCAGAGCGAGGTATGTAGGCGGTGCTACAGAGTTCTTGAAGTGGTGGCCTAACTACGGCTACACTAGAAGAACAGTATTTGGTATCTGCGCTCTGCTGAAGCCAGTTACCTTCGGAAAAAGAGTTGGTAGCTCTTGATCCGGCAAACAAACCACCGCTGGTAGCGGTGGTTTTTTTGTTTGCAAGCAGCAGATTACGCGCAGAAAAAAAGGATCTCAAGAAGATCCTTTGATCTTTTCTACGGGGTCTGACGCTCAGTGGAACGAAAACTCACGTTAAGGGATTTTGGTCATGAGATTATCAAAAAGGATCTTCACCTAGATCCTTTTAAATTAAAAATGAAGTTTTAAATCAATCTAAAGTATATATGAGTAAACTTGGTCTGACAGTTACCAATGCTTAATCAGTGAGGCACCTATCTCAGCGATCTGTCTATTTCGTTCATCCATAGTTGCCTGACTCCCCGTCGTGTAGATAACTACGATACGGGAGGGCTTACCATCTGGCCCCAGTGCTGCAATGATACCGCGAGACCCACGCTCACCGGCTCCAGATTTATCAGCAATAAACCAGCCAGCCGGAAGGGCCGAGCGCAGAAGTGGTCCTGCAACTTTATCCGCCTCCATCCAGTCTATTAATTGTTGCCGGGAAGCTAGAGTAAGTAGTTCGCCAGTTAATAGTTTGCGCAACGTTGTTGCCATTGCTACAGGCATCGTGGTGTCACGCTCGTCGTTTGGTATGGCTTCATTCAGCTCCGGTTCCCAACGATCAAGGCGAGTTACATGATCCCCCATGTTGTGCAAAAAAGCGGTTAGCTCCTTCGGTCCTCCGATCGTTGTCAGAAGTAAGTTGGCCGCAGTGTTATCACTCATGGTTATGGCAGCACTGCATAATTCTCTTACTGTCATGCCATCCGTAAGATGCTTTTCTGTGACTGGTGAGTACTCAACCAAGTCATTCTGAGAATAGTGTATGCGGCGACCGAGTTGCTCTTGCCCGGCGTCAATACGGGATAATACCGCGCCACATAGCAGAACTTTAAAAGTGCTCATCATTGGAAAACGTTCTTCGGGGCGAAAACTCTCAAGGATCTTACCGCTGTTGAGATCCAGTTCGATGTAACCCACTCGTGCACCCAACTGATCTTCAGCATCTTTTACTTTCACCAGCGTTTCTGGGTGAGCAAAAACAGGAAGGCAAAATGCCGCAAAAAAGGGAATAAGGGCGACACGGAAATGTTGAATACTCATACTCTTCCTTTTTCAATATTATTGAAGCATTTATCAGGGTTATTGTCTCATGAGCGGATACATATTTGAATGTATTTAGAAAAATAAACAAATAGGGGTTCCGCGCACATTTCCCCGAAAAGTGCCACCTGACGTC

#### Humanized E protein expression plasmid sequence

GACGGATCGGGAGATCTCCCGATCCCCTATGGTGCACTCTCAGTACAATCTGCTCTGATGCCGCATAGTTAAGCCAGTATCTGCTCCCTGCTTGTGTGTTGGAGGTCGCTGAGTAGTGCGCGAGCAAAATTTAAGCTACAACAAGGCAAGGCTTGACCGACAATTGCATGAAGAATCTGCTTAGGGTTAGGCGTTTTGCGCTGCTTCGCGATGTACGGGCCAGATATACGCGTTGACATTGATTATTGACTAGTTATTAATAGTAATCAATTACGGGGTCATTAGTTCATAGCCCATATATGGAGTTCCGCGTTACATAACTTACGGTAAATGGCCCGCCTGGCTGACCGCCCAACGACCCCCGCCCATTGACGTCAATAATGACGTATGTTCCCATAGTAACGCCAATAGGGACTTTCCATTGACGTCAATGGGTGGAGTATTTACGGTAAACTGCCCACTTGGCAGTACATCAAGTGTATCATATGCCAAGTACGCCCCCTATTGACGTCAATGACGGTAAATGGCCCGCCTGGCATTATGCCCAGTACATGACCTTATGGGACTTTCCTACTTGGCAGTACATCTACGTATTAGTCATCGCTATTACCATGGTGATGCGGTTTTGGCAGTACATCAATGGGCGTGGATAGCGGTTTGACTCACGGGGATTTCCAAGTCTCCACCCCATTGACGTCAATGGGAGTTTGTTTTGGCACCAAAATCAACGGGACTTTCCAAAATGTCGTAACAACTCCGCCCCATTGACGCAAATGGGCGGTAGGCGTGTACGGTGGGAGGTCTATATAAGCAGAGCTCTCTGGCTAACTAGAGAACCCACTGCTTACTGGCTTATCGAAATTAATACGACTCACTATAGGGAGACCCAAGCTGGCTAGCGCAGCATCTTCAGCTCTTGGGTTTAGCAAAAGTTCATCTCCTTCTGCATCCTTAACTGAGAATGAGCTTTTGTGGGAGCCCACACCAGTCAAGTTGGATTTGAACCCAGCTGCTCTGTACAAGCTTGGTACCGCCACCATGTACAGCTTCGTGAGCGAGGAGACCGGCACCCTGATCGTGAACAGCGTGCTGCTGTTCCTGGCCTTCGTGGTGTTCCTGCTGGTGACCCTGGCCATCCTG ACCGCCCTGAGGCTGTGCGCCTACTGCTGCAACATCGTGAACGTGAGCCTGGTGAAGCCCAGCTTCTACGTGTACAGCAGGGTGAAGAACCTGAACAGCAGCAGGGTGCCCGACCTGCTGGTGTAATCTAGAGGGCCCGTTTAAACCCGCTGATCAGCCTCGACTGTGCCTTCTAGTTGCCAGCCATCTGTTGTTTGCCCCTCCCCCGTGCCTTCCTTGACCCTGGAAGGTGCCACTCCCACTGTCCTTTCCTAATAAAATGAGGAAATTGCATCGCATTGTCTGAGTAGGTGTCATTCTATTCTGGGGGGTGGGGTGGGGCAGGACAGCAAGGGGGAGGATTGGGAAGACAATAGCAGGCATGCTGGGGATGCGGTGGGCTCTATGGCTTCTGAGGCGGAAAGAACCAGCTGGGGCTCTAGGGGGTATCCCCACGCGCCCTGTAGCGGCGCATTAAGCGCGGCGGGTGTGGTGGTTACGCGCAGCGTGACCGCTACACTTGCCAGCGCCCTAGCGCCCGCTCCTTTCGCTTTCTTCCCTTCCTTTCTCGCCACGTTCGCCGGCTTTCCCCGTCAAGCTCTAAATCGGGGGCTCCCTTTAGGGTTCCGATTTAGTGCTTTACGGCACCTCGACCCCAAAAAACTTGATTAGGGTGATGGTTCACGTAGTGGGCCATCGCCCTGATAGACGGTTTTTCGCCCTTTGACGTTGGAGTCCACGTTCTTTAATAGTGGACTCTTGTTCCAAACTGGAACAACACTCAACCCTATCTCGGTCTATTCTTTTGATTTATAAGGGATTTTGCCGATTTCGGCCTATTGGTTAAAAAATGAGCTGATTTAACAAAAATTTAACGCGAATTAATTCTGTGGAATGTGTGTCAGTTAGGGTGTGGAAAGTCCCCAGGCTCCCCAGCAGGCAGAAGTATGCAAAGCATGCATCTCAATTAGTCAGCAACCAGGTGTGGAAAGTCCCCAGGCTCCCCAGCAGGCAGAAGTATGCAAAGCATGCATCTCAATTAGTCAGCAACCATAGTCCCGCCCCTAACTCCGCCCATCCCGCCCCTAACTCCGCCCAGTTCCGCCCATTCTCCGCCCCATGGCTGACTAATTTTTTTTATTTATGCAGAGGCCGAGGCCGCCTCTGCCTCTGAGCTATTCCAGAAGTAGTGAGGAGGCTTTTTTGGAGGCCTAGGCTTTTGCAAAAAGCTCCCGGGAGCTTGTATATCCATTTTCGGATCTGATCAAGAGACAGGATGAGGATCGTTTCGCATGATTGAACAAGATGGATTGCACGCAGGTTCTCCGGCCGCTTGGGTGGAGAGGCTATTCGGCTATGACTGGGCACAACAGACAATCGGCTGCTCTGATGCCGCCGTGTTCCGGCTGTCAGCGCAGGGGCGCCCGGTTCTTTTTGTCAAGACCGACCTGTCCGGTGCCCTGAATGAACTGCAGGACGAGGCAGCGCGGCTATCGTGGCTGGCCACGACGGGCGTTCCTTGCGCAGCTGTGCTCGACGTTGTCACTGAAGCGGGAAGGGACTGGCTGCTATTGGGCGAAGTGCCGGGGCAGGATCTCCTGTCATCTCACCTTGCTCCTGCCGAGAAAGTATCCATCATGGCTGATGCAATGCGGCGGCTGCATACGCTTGATCCGGCTACCTGCCCATTCGACCACCAAGCGAAACATCGCATCGAGCGAGCACGTACTCGGATGGAAGCCGGTCTTGTCGATCAGGATGATCTGGACGAAGAGCATCAGGGGCTCGCGCCAGCCGAACTGTTCGCCAGGCTCAAGGCGCGCATGCCCGACGGCGAGGATCTCGTCGTGACCCATGGCGATGCCTGCTTGCCGAATATCATGGTGGAAAATGGCCGCTTTTCTGGATTCATCGACTGTGGCCGGCTGGGTGTGGCGGACCGCTATCAGGACATAGCGTTGGCTACCCGTGATATTGCTGAAGAGCTTGGCGGCGAATGGGCTGACCGCTTCCTCGTGCTTTACGGTATCGCCGCTCCCGATTCGCAGCGCATCGCCTTCTATCGCCTTCTTGACGAGTTCTTCTGAGCGGGACTCTGGGGTTCGAAATGACCGACCAAGCGACGCCCAACCTGCCATCACGAGATTTCGATTCCACCGCCGCCTTCTATGAAAGGTTGGGCTTCGGAATCGTTTTCCGGGACGCCGGCTGGATGATCCTCCAGCGCGGGGATCTCATGCTGGAGTTCTTCGCCCACCCCAACTTGTTTATTGCAGCTTATAATGGTTACAAATAAAGCAATAGCATCACAAATTTCACAAATAAAGCATTTTTTTCACTGCATTCTAGTTGTGGTTTGTCCAAACTCATCAATGTATCTTATCATGTCTGTATACCGTCGACCTCTAGCTAGAGCTTGGCGTAATCATGGTCATAGCTGTTTCCTGTGTGAAATTGTTATCCGCTCACAATTCCACACAACATACGAGCCGGAAGCATAAAGTGTAAAGCCTGGGGTGCCTAATGAGTGAGCTAACTCACATTAATTGCGTTGCGCTCACTGCCCGCTTTCCAGTCGGGAAACCTGTCGTGCCAGCTGCATTAATGAATCGGCCAACGCGCGGGGAGAGGCGGTTTGCGTATTGGGCGCTCTTCCGCTTCCTCGCTCACTGACTCGCTGCGCTCGGTCGTTCGGCTGCGGCGAGCGGTATCAGCTCACTCAAAGGCGGTAATACGGTTATCCACAGAATCAGGGGATAACGCAGGAAAGAACATGTGAGCAAAAGGCCAGCAAAAGGCCAGGAACCGTAAAAAGGCCGCGTTGCTGGCGTTTTTCCATAGGCTCCGCCCCCCTGACGAGCATCACAAAAATCGACGCTCAAGTCAGAGGTGGCGAAACCCGACAGGACTATAAAGATACCAGGCGTTTCCCCCTGGAAGCTCCCTCGTGCGCTCTCCTGTTCCGACCCTGCCGCTTACCGGATACCTGTCC GCCTTTCTCCCTTCGGGAAGCGTGGCGCTTTCTCATAGCTCACGCTGTAGGTATCTCAGTTCGGTGTAGGTCGTTCGCTCCAAGCTGGGCTGTGTGCACGAACCCCCCGTTCAGCCCGACCGCTGCGCCTTATCCGGTAACTATCGTCTTGAGTCCAACCCGGTAAGACACGACTTATCGCCACTGGCAGCAGCCACTGGTAACAGGATTAGCAGAGCGAGGTATGTAGGCGGTGCTACAGAGTTCTTGAAGTGGTGGCCTAACTACGGCTACACTAGAAGAACAGTATTTGGTATCTGCGCTCTGCTGAAGCCAGTTACCTTCGGAAAAAGAGTTGGTAGCTCTTGATCCGGCAAACAAACCACCGCTGGTAGCGGTGGTTTTTTTGTTTGCAAGCAGCAGATTACGCGCAGAAAAAAAGGATCTCAAGAAGATCCTTTGATCTTTTCTACGGGGTCTGACGCTCAGTGGAACGAAAACTCACGTTAAGGGATTTTGGTCATGAGATTATCAAAAAGGATCTTCACCTAGATCCTTTTAAATTAAAAATGAAGTTTTAAATCAATCTAAAGTATATATGAGTAAACTTGGTCTGACAGTTACCAATGCTTAATCAGTGAGGCACCTATCTCAGCGATCTGTCTATTTCGTTCATCCATAGTTGCCTGACTCCCCGTCGTGTAGATAACTACGATACGGGAGGGCTTACCATCTGGCCCCAGTGCTGCAATGATACCGCGAGACCCACGCTCACCGGCTCCAGATTTATCAGCAATAAACCAGCCAGCCGGAAGGGCCGAGCGCAGAAGTGGTCCTGCAACTTTATCCGCCTCCATCCAGTCTATTAATTGTTGCCGGGAAGCTAGAGTAAGTAGTTCGCCAGTTAATAGTTTGCGCAACGTTGTTGCCATTGCTACAGGCATCGTGGTGTCACGCTCGTCGTTTGGTATGGCTTCATTCAGCTCCGGTTCCCAACGATCAAGGCGAGTTACATGATCCCCCATGTTGTGCAAAAAAGCGGTTAGCTCCTTCGGTCCTCCGATCGTTGTCAGAAGTAAGTTGGCCGCAGTGTTATCACTCATGGTTATGGCAGCACTGCATAATTCTCTTACTGTCATGCCATCCGTAAGATGCTTTTCTGTGACTGGTGAGTACTCAACCAAGTCATTCTGAGAATAGTGTATGCGGCGACCGAGTTGCTCTTGCCCGGCGTCAATACGGGATAATACCGCGCCACATAGCAGAACTTTAAAAGTGCTCATCATTGGAAAACGTTCTTCGGGGCGAAAACTCTCAAGGATCTTACCGCTGTTGAGATCCAGTTCGATGTAACCCACTCGTGCACCCAACTGATCTTCAGCATCTTTTACTTTCACCAGCGTTTCTGGGTGAGCAAAAACAGGAAGGCAAAATGCCGCAAAAAAGGGAATAAGGGCGACACGGAAATGTTGAATACTCATACTCTTCCTTTTTCAATATTATTGAAGCATTTATCAGGGTTATTGTCTCATGAGCGGATACATATTTGAATGTATTTAGAAAAATAAACAAATAGGGGTTCCGCGCACATTTCCCCGAAAAG TGCCACCTGACGTC

